# Molecular epidemiology and high mortality of Crimean-Congo hemorrhagic fever in Oman: a re-emerging infection

**DOI:** 10.1101/502641

**Authors:** Seif S Al-Abri, Roger Hewson, Hanan Al-Kindi, Idris Al-Abaidani, Amina Al-Jardani, Amal Al-Maani, Samira Almahrouqi, Barry Atkinson, Adil Al-Wahaibi, Bader Al-Rawahi, Shyam Bawikar, Nicholas J Beeching

**Affiliations:** Seif Al-Abri, Directorate General for Disease Surveillance and Control, MoH, Muscat, Oman; Clinical Sciences, Liverpool School of Tropical Medicine, Pembroke Place, Liverpool, United Kingdom; WHO Collaborating Centre for Virus Reference and Research (Special Pathogens), Public Health England – National Infection Service, Porton Down, Salisbury, United Kingdom; Faculty of Infectious and Tropical Diseases, Dept Pathogen Molecular Biology London School of Hygiene & Tropical Medicine Institute of Tropical Medicine, Dept Emerging Disease, Nagasaki University, Japan; Central Public Health Laboratory, Directorate General for Disease Surveillance and Control, MoH, Muscat, Oman; Department of Communicable Diseases, Directorate General for Disease Surveillance and Control, MoH, Muscat, Oman; Department of Infection Prevention and Control, Directorate General for Disease Surveillance and Control, MoH, Muscat, Oman; Department of Surveillance, Directorate General for Disease Surveillance and Control, MoH, Muscat, Oman; Tropical and Infectious Disease Unit, Royal Liverpool University Hospital, Prescot Street, Liverpool, United Kingdom

**Keywords:** Crimean-Congo hemorrhagic fever, CCHF, viral, Oman

## Abstract

**Background:** Crimean-Congo hemorrhagic fever (CCHF) is a serious disease with a high fatality rate reported in many countries. The first case of CCHF in Oman was detected in 1995 and serosurveys have suggested widespread infection of humans and livestock throughout the country.

**Methodology:** Cases of CCHF reported to the Ministry of Health (MoH) of Oman between 1995 and 2017 were retrospectively reviewed. Diagnosis was confirmed by serology and/or molecular tests in Oman. Stored RNA from recent cases was studied by sequencing the complete open reading frame (ORF) of the viral S segment at Public Health England, enabling phylogenetic comparisons to be made with other S segments of strains obtained from the region.

**Findings:** Of 88 cases of CCHF, 4 were sporadic in 1995 and 1996, then none occurred until 2011. From 2011-2017, incidence has steadily increased and 19 (23.8%) of 80 cases clustered around Eid Al Adha. The median (range) age was 33 (15-68) years and 79 (90%) were male. The major risk for infection was contact with animals and/or butchering in 73/88 (83%) and only one case was related to tick bites alone. Severe cases were over-represented: 64 (72.7%) had a platelet count < 50 x 10^9^/L and 32 (36.4%) died. There was no intrafamilial spread or healthcare-associated infection. The viral S segments from 11 patients presenting in 2013 and 2014 were all grouped in Asia 1 (IV) lineage.

**Conclusions:** CCHF is well-established throughout Oman, with a single strain of virus present for at least 20 years. Most patients are men involved in animal husbandry and butchery. The high mortality suggests that there is substantial under-diagnosis of milder cases. Preventive measures have been introduced to reduce risks of transmission to animal handlers and butchers and to maintain safety in healthcare settings.

**Author summary:** Crimean-Congo hemorrhagic fever, an often fatal tick-borne viral disease, has made an impact in the Sultanate of Oman—affecting nationals and expatriates alike—for the past 20 years. In this retrospective review of the epidemiology and outcomes of cases in Oman from 1995 to 2017, we identified 4 sporadic cases in 1995 and 1996, then none until 2011, followed by a steady increase until 2017. The mortality rate of 32 of 88 cases (36.4%) is high in comparison to studies from other countries and this could be explained by under-diagnoses of milder cases in the Sultanate.

Transmission is commonly associated with animal husbandry and butchering and 88% cases were infected by contact with animals, whereas transmission by tick bite is more commonly recorded in some countries. A proportion of cases (23.8%) were clustered around the Eid-Al-Ahda festival which has, from 2011-2017, occurred in the summer months, which have a higher risk of transmission. This additional risk has been noted and preventive measures have been introduced to reduce the risk of transmission to animal handlers and butchers.

## Introduction

Crimean-Congo hemorrhagic fever (CCHF) is a serious and often fatal infection caused by the CCHF virus (CCHFV). Ixodid ticks, especially *Hyalomma spp*, act as both reservoirs and vectors. This virus has the greatest geographic range of any tick-borne virus and there are reports of viral isolation and/or disease from more than 30 countries in Africa, Asia, Eastern and Southern Europe, and the Middle East [1–3]. Numerous domestic and wild animals, such as cattle, goats, sheep and small mammals, such as hares and rodents, serve as asymptomatic amplifying hosts for the virus [4].

CCHFV can be transmitted between animals and humans by *Hyalomma* ticks. It can also be transmitted by direct contact with blood and other body fluids of viremic humans and animals and has the potential to cause population-based outbreaks [5–6]. Clinical features commonly include fever of abrupt onset, myalgia, headache and thrombocytopenia, and can progress to hemorrhage, multiorgan failure and death. The levels of liver enzymes, creatinine phosphokinase, and lactate dehydrogenase are raised, and bleeding markers are prolonged [7–8]. The crude mortality rate of CCHF differs from country to country, ranging from 2-80% [1]. Early diagnosis and supportive management are essential for a favorable outcome.

CCHFV is a negative-sense single-stranded RNA virus classified within the *Orthonairovirus*genus of the *Nairoviridae* family. The CCHFV genome is comprised of single-stranded negative-sense RNA divided into 3 distinct segments designated small (S), medium (M), and large (L). Comparisons of full S segment sequences have shown that CCHFV forms 7 distinct clades, each with strong geographical associations [1,9–11]. Subtle links between distant geographic locations, shown by phylogenetic analysis, may have originated from the international livestock trade or from long-distance carriage of CCHFV by infected ticks via bird migration [5,10,12–13].

Oman is situated in the southeastern corner of the Arabian Peninsula, bordering the Kingdom of Saudi Arabia, United Arab Emirates, and Yemen. The summer is hot and humid with temperatures reaching as high as 49°C and the winter relatively cooler and with some rain. The total population is 4,615,269 individuals, of whom 54.6% are Omanis and the remainder are expatriates [14].

Cases of CCHF were first detected in Oman in 1995 when there were 3 unrelated sporadic cases, followed by a further case in 1996 [15, 16]. Cases related to animal movement and slaughter were also reported the following year from Western Saudi Arabia [17–18] and from the UAE [15,19, 20], where an imported case had previously resulted in fatal infections of health care workers in 1979 [21]. A survey conducted in 1996 in Oman revealed asymptomatic seropositivity for CCHFV exposure in 1/41 (2.4%) of Omanis compared to 73 (30.3%) of 241 non-Omani citizens with occupational animal contact [22]. However, no further human cases of CCHF were reported in Oman until 2011 [23], and since then there has been a steady increase [24]. A recent survey has shown infection in a variety of animals and ticks in Oman [25].

Limited data are available on the prevalent clade(s), or group(s) of organisms from a single ancestor, of CCHFV in the Arabian Peninsula. Sequencing of the S, M, and L segments of CCHFV isolated from the 1996 patient in Oman (recorded as Oman 1997 in GenBank) showed that it belonged to Asia lineage 1 (clade IV) [10,26], as was the virus isolated from a patient who returned to India with CCHF acquired in Oman in 2016 [27]. Virus isolates from 4 patients in the UAE in 1994 and 1995 also align with the Asia 1 (clade IV) lineage, as did contemporaneous isolates from *Hyalomma* ticks obtained from livestock imported into the UAE from Somalia [9,19,28]. A further human isolate in the UAE in 1994/95 aligned with lineage Africa 1 (clade III) [9, 19, 28].

The aims of this study are to describe the clinical and epidemiological features and outcomes of cases of CCHF diagnosed in Oman between 1995 and 2017. We also investigated the local molecular epidemiology of CCHFV by partial and complete S segment sequencing of stored CCHFV isolates from patients recently diagnosed in Oman.

## Methods

A retrospective descriptive record-based review and analysis of CCHF cases was conducted over the period 1995 through 2017. CCHF has been listed as a notifiable disease in Oman since 1995 and surveillance forms from suspected cases are submitted by all healthcare providers to the Communicable Diseases Department at MoH headquarters. Blood samples obtained from suspected cases were submitted at the same time to the Central Public Health Laboratory (CPHL) at the MoH in Muscat, Oman. All CCHF cases reviewed and included in this study were detected by this routine communicable disease surveillance combined with the CPHL results during the study period.

A generic national form is used for initial notification of a suspected case of CCHF; once the diagnosis is confirmed, a more detailed form is submitted that includes patient identifiers, demographic and geographic variables, relevant exposure history, key clinical features, and some clinical laboratory test results. The form is not unique to CCHF, so prompts for some specific CCHFV-related exposures and laboratory variables are missing. The report format remained similar until 2017 when the paper form was replaced by an electronic version.

Data were systematically extracted from the surveillance forms of all laboratory confirmed cases of CCHF. Demographic variables included age, sex, nationality, location, and date of notification. Risk factors included history of tick bite, occupational exposure and contact with tissues, blood or other biological fluids from an infected animal, or contact with a case within 14 days prior to the onset of symptoms. Clinical data included presence/absence and duration of fever, headache, myalgia, nausea, vomiting, diarrhea, petechial rash, and bleeding from sites including gums, nose, lung, gastrointestinal tract, or skin. Key laboratory variables include platelet counts, hemoglobin, urea and electrolytes, and liver function tests. The main clinical outcomes were death or survival.

The case definition for a suspected case in Oman is: an illness with sudden acute onset with the following clinical findings: a fever ≥ 38.5°C (> 72 hours to < 10 days) associated with severe headache, myalgia, nausea, vomiting, and/or diarrhea; thrombocytopenia < 50 x 10^9^/L; hemorrhagic manifestations which develop later and may include petechial rashes, bleeding from the gums, nose, lungs, gastrointestinal tract, etc.; history of tick bite, occupational exposure, contact with fresh tissues, blood, or other biological fluids from an infected animal [24].

Good laboratory practice and a high level of effective biosafety precautions are required by laboratory staff handling materials from suspected CCHF cases due to the potential for sample-to-person, or indirect, transmission [6,29]. Blood samples collected from suspected cases of CCHF admitted to all MoH and non-MoH health care institutions in Oman are sent to CPHL in triple pack containers, using the most direct and timely route available. These samples are considered urgent and results are provided within 24 hours of their arrival at CPHL. National guidelines are in place to instruct local laboratories where suspected cases are admitted on safe handling of all material collected for any diagnostic purpose [24,30].

Both serum and plasma samples are requested for CCHFV testing. Plasma is preferred for molecular testing using a commercial CCHFV real time reverse transcription polymerase chain reaction (rRT-PCR) kit (In vitro Diagnostics, Liferiver Shanghai ZJ Bio-Tech Co., Ltd. Shanghai, China). At CPHL, the plasma extraction takes place inside a gloved box using a manual extraction system, QIAamp Viral RNA Kit (QIAGEN, Hilden, Germany). The samples are first treated with AVL buffer (QIAGEN) to inactivate infectious viruses and RNases. Intact viral RNA is then purified by selective binding and washing steps. The screening RT-PCR reaction is based on a one step real-time RT-PCR. Briefly, CCHFV RNA is converted into cDNA and a thermostable DNA polymerase is used to amplify specific CCHFV S segment sequence targets by standard thermocycling in a PCR as per manufacturer instruction. The kit contains an internal control to identify possible PCR inhibition.

A positive result from a RT-PCR screening for CCHFV RNA is used to confirm infection. In such cases, the serum sample is not tested further. If the RT-PCR is negative, heat inactivated serum (56°C water bath for 30 minutes) is tested for CCHFV antigen and IgM and IgG antibodies using a commercial kit (Vector-Best, Novosibirsk, Russia). For samples that are negative for all parameters, a convalescent serum is requested for CCHF IgG testing. The CPHL takes part in regular internal and external quality assurance reviews in association with WHO EMRO and WHO Quality Management Standards.

All available stored serum samples, collected from 21 CCHF patients in 2013 and to 2014, were inactivated with AVL buffer and sent to PHE Porton Down, England, UK. At PHE, AVL samples were processed with a standard QIAamp Viral RNA Kit. Eluted RNA was evaluated for the presence of CCHFV RNA using an in-house RT-PCR assay [31]. Sequencing was performed using standard CCHFV S segment sequencing primers as described previously [32]. Assembled sequence data for the S segment of each sample were manipulated and analyzed using the Lasergene suite of programs (DNAStar, Maddison, WI, USA). For phylogenetic analysis, sequences were aligned using the Clustal W computer program (The European Bioinformatics Institute, Wellcome, UK) [33] and output in PHYLIP Format (scikit-bio). To construct maximum-likelihood phylogenetic trees, quartet puzzling was applied using the program, Tree-Puzzle, at the Institut Pasteur [34, 35]. The Tamura-Nei model of substitution was adopted, as has been performed in other phylogenetic studies demonstrating reassortment [36]. Phylogenetic trees were drawn using the program TreeView (JAM Software GmbH, Trier, Germany) [37]. The values at the tree branches represent the puzzle support values. S segment sequences were submitted to GenBank.

The data analysis was conducted at the MoH Department of Surveillance in Muscat. A descriptive analysis compared age, sex, nationality, location, and date of cases. Risk factors and clinical and laboratory parameters were also tabulated. Missing data items (positive or negative) were omitted from analysis. Statistical comparisons were performed using SPSS 11.0 package program (SPSS Inc, Chicago, IL, USA).

Ethical approval was sought from the MoH, Oman. The study is considered free from ethical constraints as it is a secondary analysis of the data collected routinely for the purpose of public health surveillance and reporting. No personal identifying information accompanied the samples sent to PHE.

## Results

A total of 88 cases were reported between 1995 and 2017. Of these, 82 (93.2%) were confirmed by RT-PCR and 4 by CCHFV IgM alone. Two further probable cases (both fatal) in 2011 and 2016 were included on the basis of typical clinical and laboratory features as per electronic records. There were 3 isolated cases in January, May, and June 1995 with a further case in 1996, and then no cases were reported until 2011. Since then, there has been a steady increase in numbers, peaking at 20 cases in 2015 (Fig 1). Annual notifications of suspected cases also dipped between 1996 and 2011 (Fig 1). Annual notifications of suspected cases were not systematically recorded until 2011 and data about notifications of suspected cases and possible missed cases are incomplete. In the years 2001 to 2011 inclusive, there were 35 notifications of possible VHF cases, of which 2 were proven CCHF (in 2011) and at least 23 were confirmed to be cases of dengue (2 fatal).

**Fig 1.**
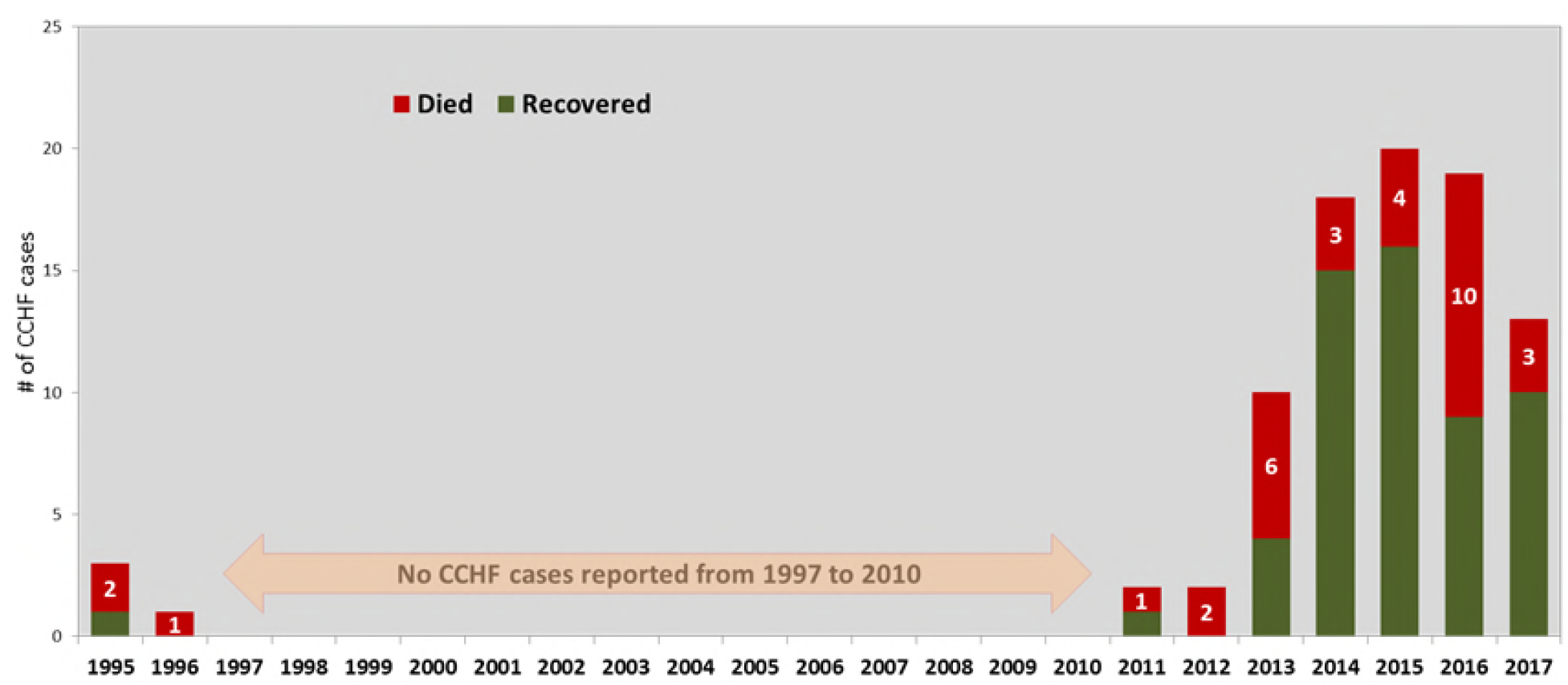
Yearly distribution of CCHF cases and deaths reported in Oman: 1995-2017.

The patients had a median (range) age of 33 (15-68) years and 79 (90%) were male. The most common nationality affected was Omani 51 (59%) followed by Bangladeshi 18 (21%), Pakistani 7 (8%), Yemeni 3 (4%), Indian 4 (5%), Somali 2, and Sri Lankan 1. Cases occurred in all governorates (wilayats) except Musandam and Al Wustah (Fig 2). There was no geographic or source-related clustering of cases; however, several cases followed Eid Al Adha, a festival associated with animal sacrifice. In the years 2013-2017, 19/80 (23.8%) of all cumulative cases had their onset within 3 weeks after Eid Al Adha (Fig 3). There was also a smaller peak of cases in the spring weeks 6-19 (Fig 3). The main exposure risk identified was animal/fresh tissue exposure in 73/88 (83%), with only 1 case attributed to tick bite alone. Exposure risk was not identified in 14 (15.9%) (Table 1).

**Fig 2.**
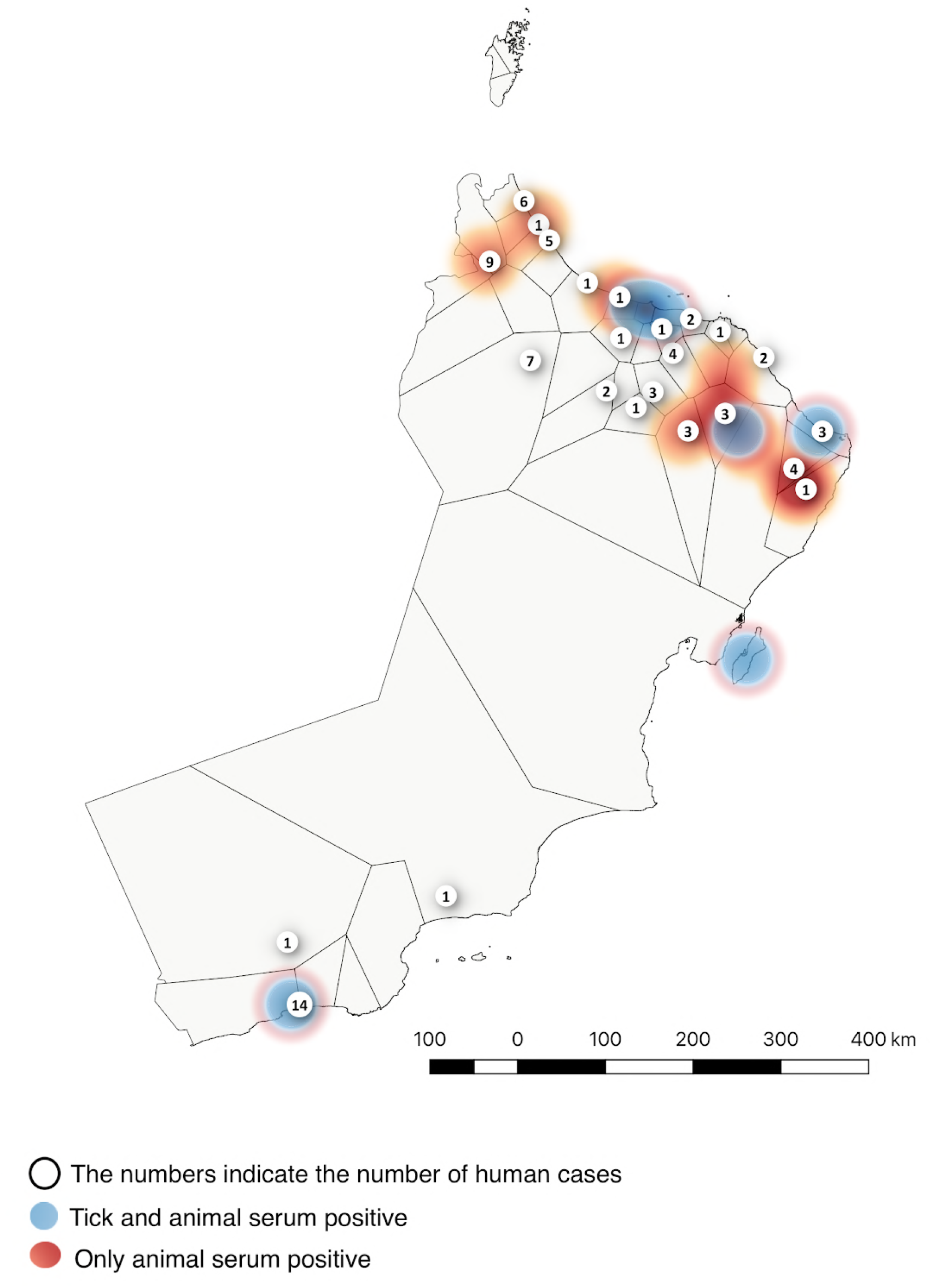
Map of Oman with number of CCHF cases, superimposed on results of previous animal serosurveys for CCHFV antibodies and tick surveys for CCHFV (24-25).

**Fig 3.**
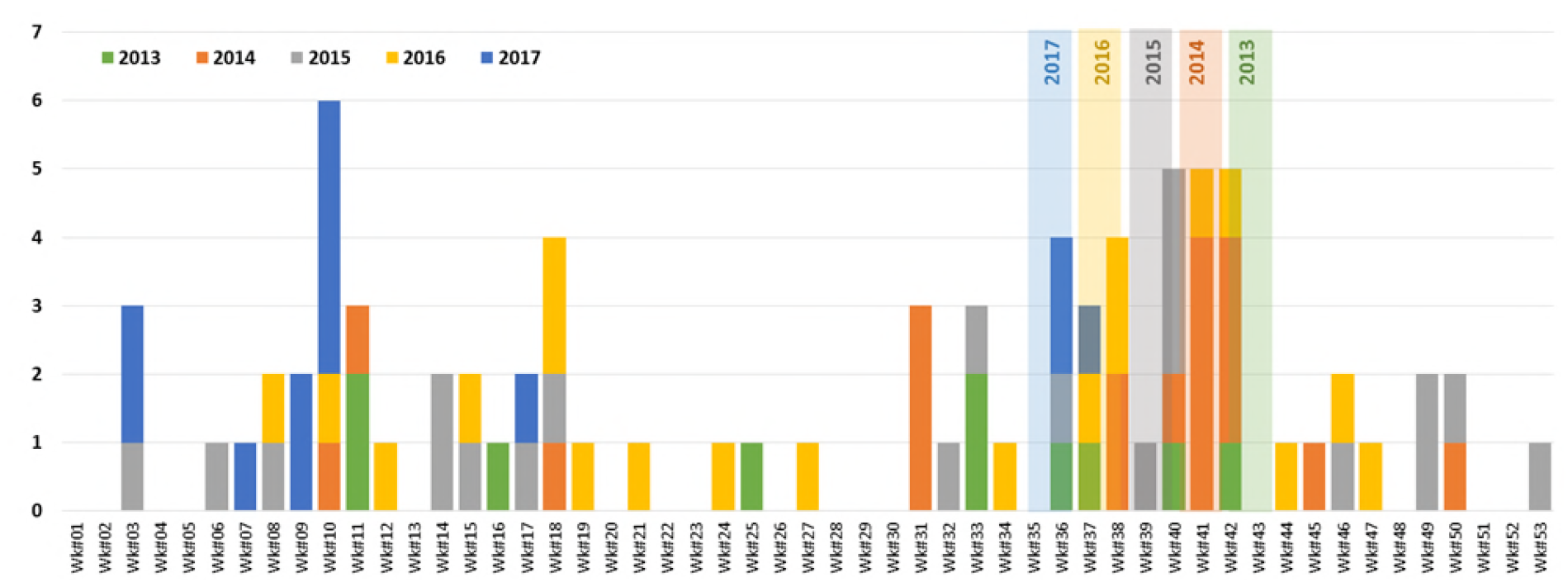
Weekly distribution of CCHF cases in Oman showing clustering after Eid Al Adha: 2013 – 2017 (n=80.

Eid Al Adha occurs 10 days earlier in each successive year, indicated by the pale vertical columns between weeks 35 and 43. The risk period is defined as approximately 10 days after the Eid Al Adha festival.

**Table 1.**
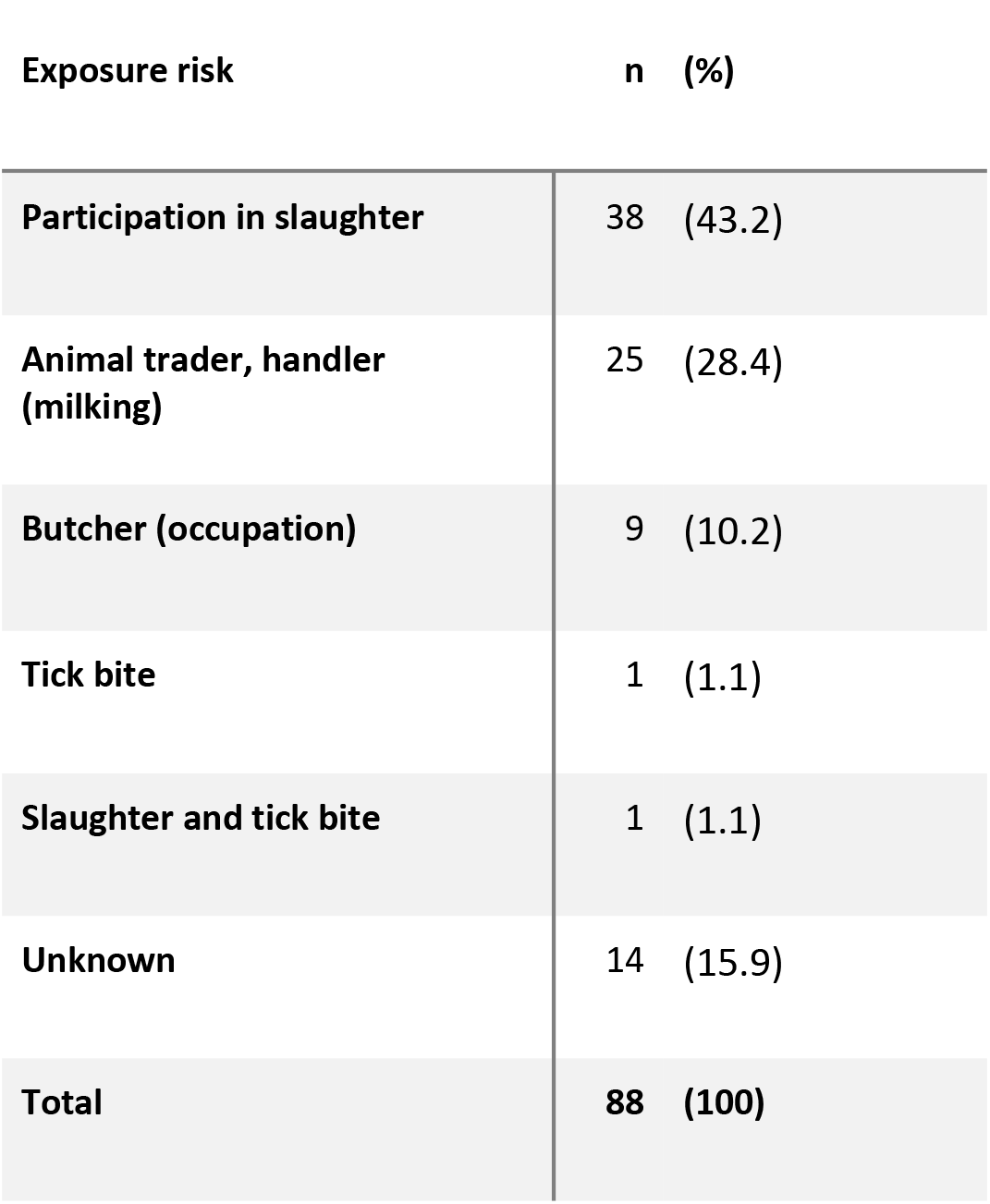
Exposure risk factors for all 88 patients.

| **Exposure risk** | **n (%)** |
| --- | --- |
| **Participation in slaughter** | **38 (43.2)** |
| **Animal trader, handler (milking)** | **25 (28.4)** |
| **Butcher (occupation)** | **9 (10.2)** |
| **Tick bite** | **1 (1.1)** |
| **Slaughter and tick bite** | **1 (1.1)** |
| **Unknown** | **14 (15.9)** |
| **Total** | **88 (100)** |

Clinical features in 88 patients included fever in 80 (90.9%), hemorrhagic features 41 (46.6%), vomiting 32 (36.4%), myalgia 30 (34.1%), diarrhea 20 (22.7%), respiratory symptoms 17 (19%), abdominal pain 11 (12.5%), other symptoms in 29 (33%). Severe thrombocytopenia (platelet count < 50 x 10^9^/L) was present in 64 (72.7%). There were 32 deaths, resulting in a cumulative case fatality rate of 36.4%. The case fatality rate in Omanis was 16/53 (30.2%) and in Bangladeshis was 10/18 (55.6%) (P>0.05).

Of the 21 serum samples that were sent to PHE, 20 were RT-PCR positive using an in-house assay. However, of these, only 12 samples provided suitable cycle threshold values (the cycle threshold being 28 or under) to warrant further sequencing of CCHFV S segments and only 12 samples provided sequencing data which spanned the entire ORF of the S segment. Sequence data have been submitted to GenBank and sequences have been assigned the following accession numbers: MH037279 (Oman 2012-40S), MH037280 (Oman 2013-116S), MH037281 (Oman 2014-828P), MH037282 (Oman 2014-979P), MH037283 (Oman 2014-602P), MH037284 (Oman 2013-825P), MH037285 (Oman 2013-92S), MH037286 (Oman 2013-108S), MH037287 (Oman 2013-179P), MH037288 (Oman 2014-860P), MH037289 (Oman 2014-624S), and MH037290 (Oman 2014-747P). Sequences were compiled with a range of other CCHFV S segment ORF sequences and used to make the maximum likelihood phylogenetic tree shown in Figure 4.

**Fig 4.**
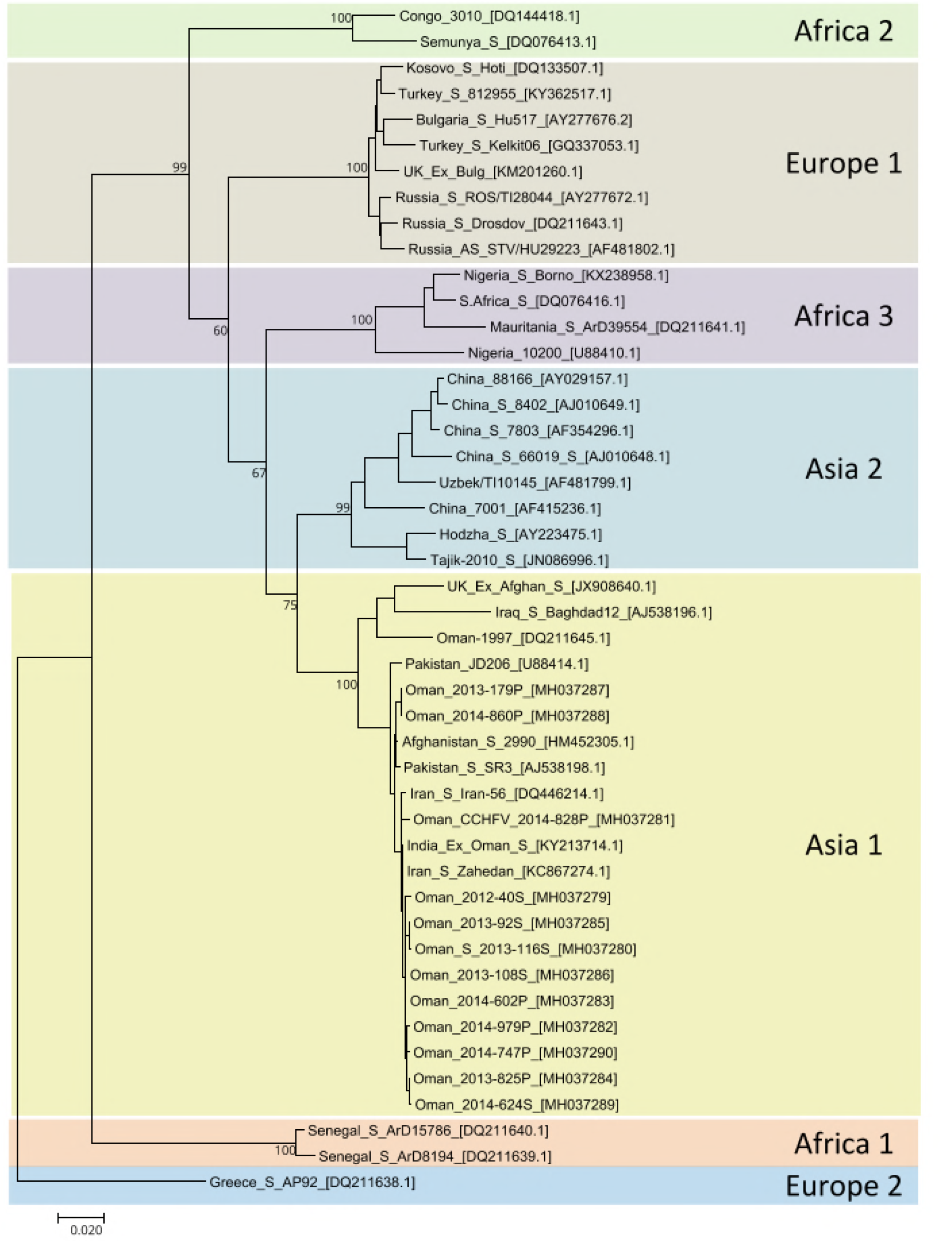
Phylogenetic Tree showing Omani sequences clustering in Asia 1 lineage, with other sequences from the GCC region previously deposited in Genbank.

## Discussion

This report summarizes the clinical, epidemiological, and virological findings in 88 people with symptomatic CCHF throughout the Sultanate of Oman in the past 2 decades. Cases were detected by passive surveillance, starting with a few sporadic reports in 1995 and 1996, followed by no cases until 2011. Since then there has been a sustained increase in yearly reports, of which 19/80 (23.8%) have clustered around the Eid Al Adha festival, occurring in summer months in the years 2013 to 2017, with a possible smaller peak in the spring months. Ninety percent of all patients were male with a median age of 33 years. Both Omanis and citizens of other nationalities were affected, the predominant risk factors being exposure to animals and meat products, especially involvement in butchering or slaughtering. Diagnosis was confirmed by RT-PCR in 82 (93.2%) cases and by serology alone in 4 (4.5%). Stored viral RNA from 12 patients presenting in 2013 and 2014 was sequenced for the entire S segment ORF of each of the 12 samples, all grouped in the Asia 1 (IV) clade. The cumulative mortality was 36.4%, and no cases of healthcare related or intrafamilial spread of infection were reported.

This is the largest series of cases of CCHF reported from a GCC country, and provides the first data about locally prevalent strains of CCHFV in almost 20 years. The findings raise a number of questions about the origin and distribution of CCHFV in Oman and neighboring countries, the reasons for the high observed mortality, and the appropriate human and veterinary public health responses in Oman and other GCC states.

The clinical features of the cases were similar to those reported in other countries [13,38–40].The most common symptoms reported were fever, fatigue, headache, loss of appetite, myalgia, and abdominal pain. Hemorrhagic manifestations were described in 34/67 (50.8%) and severe thrombocytopenia (< 50 x10^9^/L) was present in 64/88 (72.7%) at presentation. There is no internationally agreed case definition for CCHF, but at least 3 scoring systems to assess severity of illness have been proposed [41–43]. Mortality is known to be associated with older age, presence of underlying illness, and CCHFV viral load at presentation [8]. Case numbers were too small to show a link with mortality in our series and details about the latter 2 risks were not recorded. Representative case fatality rates elsewhere include 5% in Turkey, 17.6% in Iran, and 15% in Pakistan [1,44–45]. However, the CFR of 15% in Pakistan was reported from a center with substantial experience, whereas overall mortality rates of up to 41% have been reported more recently in Pakistan, especially during outbreaks [46].

The lower mortality in Turkey and Iran could be explained by improved surveillance and early diagnosis of CCHF in patients with fever and thrombocytopenia, following prolonged campaigns to raise awareness in both healthcare personnel and the general public in those countries. A serosurveillance study conducted in Oman in 1996 showed that none of the 74 antibody-positive individuals identified recalled ever being hospitalized for an illness resembling CCHF with associated fever and bleeding, suggesting that there is a substantial incidence of subclinical CCHF human infections in Oman [3, 12–13,22]. Serosurveillance studies in other countries have shown seroprevalence rates of approximately 10-13% in high risk human populations [5, 47]. Based on mortality data from Turkey we believe that under-diagnosis of mild cases has skewed the mortality data in Oman.

This is the first study to describe the complete sequence of the S segment ORFs of a series of CCHFV isolates from the region. The results largely confirm findings from partial sequencing of sporadic isolates from the UAE [19, 28] and Oman [10,27] since the mid 1990’s. The phylogenetic relationship of these sequences with other published sequences from the region is depicted in Figure 4. The similarity of all these sequences to the human cases in Oman and the UAE over 2 decades is striking, and these sequences align with those from Pakistan. In the future, with the advance of cheaper sequencing technologies, it will be valuable to compare full length genomes from multiple locations with clinical data. This may help address hypotheses about alternative strain pathogenicity, including the relative contribution of segment reassortment in CCHF disease [10, 48].

The average age of patients in Oman was 33 years, and 90% were men. Most infections were acquired while butchering or slaughtering animals or from other close animal tissue and blood exposure as in earlier and more recent cases in Dubai [20,49] and the Kingdom of Saudi Arabia [17]. This contrasts with the situation in Turkey, Kazakhstan, and Iran, where tick bites are the most commonly reported risk factor [11, 39–40,44]. Slaughtering animals during Eid Al Adha is known to pose a particularly high risk of infection [5]. Sporadic unregulated slaughtering without using appropriate personal protective equipment still occurs in Oman during Eid Al Adha. It is also common for non-professional individuals to become involved and for butchers to freelance, going from house to house to sacrifice animals, as people find it more convenient to have the sacrifice performed at farms and backyards. During skinning and subsequent tanning of the hides, ticks can bite humans. We examined the effect of Eid as a possible cause for the apparent increase in cases over the past 7 years as the festival has moved back into the summer months when tick activity is most prominent. However, this is not the only factor. Figure 3 demonstrates that there is the expected clustering of cases after Eid Al Adha, but this accounts for only 23.6% of the total cases. Similar findings have been reported in Pakistan [50] and the data suggest that climatic factors affecting tick activity are most important in promoting seasonal variation in human infection risk together with extra added risk at the time of Eid.

These data change our perceptions about the duration and origin of CCHFV activity in Oman and neighboring countries. Previously, it had been postulated that sporadic cases were related to the importation of infected livestock from other countries. A large amount of livestock is imported to Oman every year: in 2016, over 1.5 million farm animals were imported, including sheep (85.2%), cattle (7%), and goats (5.6%). The origins of these included Armenia, Australia, Djibouti, India, Iran, Jordan, Pakistan, Somalia, Sudan, Turkey and the UAE. A 21-day quarantine procedure is in effect for animals arriving from other countries by sea or land. Once in Oman, animals are distributed to sales centers, feedlots, and distribution points throughout the country. Livestock are held in large holding pens and not segregated according to country of origin or time from entry into Oman. Spread of infection could result from unrestricted entry of tick-infested and potentially viremic domestic animals during religious holidays; the abundance of virus-infected ticks within stockyards and holding pens; the uncontrolled movement of livestock animals infested with CCHFV-carrying *Hyalomma* ticks to ranches, farms, and markets throughout the country; and the indiscriminate mixing and crowding of tick-infested and potentially viremic animals with uninfected and tick-free animals [3,13,22].

There was partial support for the possibility of intermittent importation of CCHFV with livestock into the UAE in the 1990s, where ticks were found on animals with different clades of CCHFV S segment corresponding to African as well as Asia 1 clades [9,19,28]. However, human serosurveys in Oman in 1996 [22] and the finding of the virus in ticks and animals throughout Oman in 2013-2014 [25] suggest that all areas of the Sultanate have had a substantial burden of CCHFV infection for at least 2 decades, probably related to all the risks mentioned above. Moreover, all virus isolates from humans in Oman and the UAE have had remarkably similar S Segments, apart from the nosocomial outbreak in Dubai in 1979 from an Indian index case [21]. In contrast, several different S segment strains of virus are circulating in Somalia, Iran, and Turkey [9,10,51]. This suggests that the Asia 1 S segment of CCHFV has been in endemic in Oman for more than 20 years. It will be of interest to fully sequence the complete genomes of the recent isolates from ticks in Oman (and elsewhere) to explore this hypothesis further [25]. Reports of CCHFV antibody positivity in earlier human serosurveys in Kuwait [52] and intermittent occupational-related outbreaks in the UAE [19, 20, 28], and KSA [17,18], since then suggest that this is also the case throughout GCC countries.

The Oman MoH has undertaken a number of activities and initiatives to educate and inform the public about the risks of CCHF infection associated with slaughtering. A joint strategic initiative was developed in collaboration with the Ministry of Agriculture and Fisheries and Ministry of Municipalities and Water Resources. Education and information on prevention of CCHF in different languages has been targeted at those involved in slaughtering and handling animals. This includes placing advertisements on social media platforms, TV, radio, billboards, magazines, and newspapers before and during Eid Al Adha. Knowledge about CCHF is increasing in Oman with hospitals now following guidelines for the management of suspected cases of CCHF [53]. In addition, guidelines have been produced for culturally acceptable safe burials [24, 54]. It is reassuring that no healthcare related infections were detected in this series.

The data suffer from the limitations of a retrospective study that spans over 20 years, based on notifications of suspected illness and laboratory reports. In particular, the completeness of notification has been highly variable and is likely to have underestimated the incidence of symptomatic infections. Data on notifications of suspected cases that later turned out to be negative for CCHFV have not been systematically recorded and it is possible that the gap in observed cases between 1997 and 2011 is due to underreporting or referral. However, the records of confirmed cases at CPHL are thought to be complete. This is the largest reported series of CCHF from any of the GCC countries to date and brings together all published viral sequences in this region. The implication is that CCHF is endemic and under-recognized in Oman and surrounding countries and that prospective studies are needed to determine how often less severe cases of fever and thrombocytopenia are presenting in Oman. Proven and suspected cases have been reported in expatriate travelers returning from Oman to India [27] and Pakistan [55] and the possibility of CCHF should be considered in febrile travelers arriving from GCC countries, especially if they have been involved in animal slaughtering [56]. Oman has responded by improving its notification systems and laboratory support. Active local and regional programs of health promotion and human illness prevention need to be maintained together with surveillance and control of infection in animals and local tick vectors.

## Acknowledgments

The authors are pleased to acknowledge the staff in the Central Public Health Laboratory (Muscat), the Department of Communicable Diseases, the Department of Surveillance, and the Department of Infection Prevention and Control at the Directorate General for Disease Surveillance and Control (Muscat), and the staff in PHE in Porton Down (UK) for their great efforts in supporting the conduct of the study. RH and NJB are affiliated to the National Institute for Health Protection Research Unit (NIHR HPRU) in Emerging and Zoonotic Infections at University of Liverpool in partnership with PHE, in collaboration with the Liverpool School of Tropical Medicine. RH is based at PHE, Porton and NJB is based at the Liverpool School of Tropical Medicine. The views expressed are those of the author(s) and not necessarily those of the NHS, the NIHR, the UK Department of Health or PHE.

**Supporting Information Legends:** S1 STROBE Checklist

